# Altered *N*-glycan composition impacts flagella mediated adhesion in *Chlamydomonas reinhardtii*

**DOI:** 10.1101/2020.05.18.102624

**Authors:** Nannan Xu, Anne Oltmanns, Longsheng Zhao, Antoine Girot, Marzieh Karimi, Lara Hoepfner, Simon Kelterborn, Martin Scholz, Julia Beißel, Peter Hegemann, Oliver Bäumchen, Luning N. Liu, Kaiyao Huang, Michael Hippler

## Abstract

For the unicellular alga *Chlamydomonas reinhardtii*, the presence of *N*-glycosylated proteins on the surface of two flagella is crucial for both cell-cell interaction during mating and flagellar surface adhesion. It is unknown whether the composition of *N*-glycans attached to respective proteins is important for these processes. To this end, we examined several *C. reinhardtii* insertional mutants and a CRIPSR/Cas9 knockout mutant of xylosyltransferase 1A, all possessing altered *N*-glycan compositions. Taking advantage of atomic force microscopy and micropipette force measurements, our data revealed that reduction in *N*-glycan complexity impedes the adhesion force required for binding the flagella to surfaces. In addition, polystyrene bead binding and transport is impaired. Notably, assembly, Intraflagellar Transport and FMG-1B transport into flagella are not affected by altered *N*-glycosylation. Thus, we conclude that proper *N*-glycosylation of flagellar proteins is crucial for adhering *C. reinhardtii* cells onto surfaces, indicating that *N*-glycans mediate surface adhesion via direct surface contact.

## Introduction

*N*-glycosylation, as one of the major post-translational modifications, takes place along the ER/Golgi secretion route and consequently most *N*-linked glycans are found on proteins facing the extracellular space. Initial steps of *N*-glycosylation in the ER are highly conserved among most eukaryotes and consist of the synthesis of a common prebuilt *N*-glycan precursor onto a dolichol phosphate. Following the transfer of the glycan precursor onto the asparagine of the consensus sequence N-X-S/T of a nascent protein (where X can be any amino acid except proline), the glycoprotein is folded by the glycan recognizing chaperones Calnexin and Calreticulin (Stanley P, Taniguchi N, 2017). Subsequent *N*-glycan maturation steps in the Golgi are species dependent and gives rise to a high variety of *N*-glycan structures. In land plants, Golgi maturation leads to *N*-glycans modified with β1,2-core xylose and α1,3-core fucose (Strasser, 2016). In *Chlamydomonas reinhardtii*, a unicellular biflagellate green alga, *N*-glycans can be decorated with core xylose and -fucose (Lucas et al., 2020; Oltmanns et al., 2020; Schulze et al., 2018). Additionally, *6O*-methylation of mannose and addition of a terminally linked β1,4-xylose were reported (Mathieu-Rivet et al., 2013). While the functional advantage of *N*-linked glycans of mature proteins is hardly understood, it is known that blocking the synthesis of a full *N*-glycan precursor results in hypoglycosylated proteins that cannot be folded properly (Gardner et al., 2013). Instead, they are degraded via the ER-associated degradation pathway (ERAD) and are consequently not targeted correctly (Adams et al., 2019; Cherepanova et al., 2016). Therefore, impairment of glycosylation at such early stage is lethal in both uni-and multicellular organisms and only when inhibition of glycosylation is carefully dosed (e.g. by tunicamycin), immediate physiological effects caused by hypoglycosylation can be observed (Kukuruzinska et al., 1987). Looking at *C. reinhardtii*, treatment of vegetative cells with tunicamycin lead to an impaired flagellar adhesiveness, indicating that glycoproteins are crucial for adhesion and the subsequent onset of gliding (Bloodgood, 1987). However, whether this phenotype is linked to lack of proteins and/or the of lack *N*-glycosylation is unclear.

Whole cell gliding is one of two flagella based motilities in *C. reinhardtii* besides swimming (Ishikawa and Marshall, 2011; Kozminski et al., 1993; Snell et al., 2004). In principle, the cell adheres to a surface via its flagella, positioning them in a 180° angle and starts gliding along the solid or semisolid surface into the direction in which one flagellum is pointing (designating it as leading flagellum) (Bloodgood, 2009). Interestingly, flagella not only bind to large solid surfaces, but they also bind to small, inert objects (e.g. polystyrene microbeads) that are moved along the flagellar membrane. While the two events are believed to underly the same molecular machinery, it is assumed that they start with an adhesion of flagella membrane components to the surface (Bloodgood and Salomonsky, 1998). A micropipette force measurement approach recently showed that the flagella adhesion forces on different model surfaces with tailored properties lie in the range of 1 to 4 nN and that only positive surface charge diminished the adhesion force significantly (Backholm and Bäumchen, 2019; Kreis et al., 2019, 2018). Remarkably, surface iodination experiments in the early 1980s revealed a single protein called flagellar membrane glycoprotein 1B (FMG-1B) as the main player mediating surface contact (Bloodgood and Workman, 1984). FMG-1B is exclusively located in the flagellar membrane and has a remarkable size of around 350 kDa (4389 amino acids) with a large extra-flagellar part (4340 amino acids) anchored in the membrane via an expected single trans membrane helix of 22 amino acids (Bloodgood et al., 2019). As the name indicates, it is heavily *N*- and *O*-glycosylated. A recent knock down study showed, that it is the main constituent of the glycocalyx surrounding the flagellum. Additionally, a *fmg-1B* mutant showed a drastically reduced ability to glide (Bloodgood et al., 2019). Strikingly, FMG-1B is present at a high copy number and turns over rapidly within approximately 1 h (Bloodgood, 2009). The rapid turnover is probably attributed to the fact that flagellar membrane components are constantly shed into the medium as flagellar ectosomes (Bloodgood, 2009; Wood et al., 2013). FMG-1B and another *N*-glycosylated membrane component, FAP113, have been shown to be eventually torn out of the membrane once bound to microbeads (Kamiya et al., 2018). Whether FAP113 is only involved in microbead binding or also in whole cell gliding is unknown.

Recently, it was found that the flagella adhesion to surfaces is switchable by light, indicating that a blue-light photoreceptor signal is underlying this process.^18^ Following adhesion to the surface, a transmembrane signal mediates translation of the adhesion event into a calcium transient and protein phosphorylation cascade (Bloodgood, 2009; Collingridge et al., 2013; Kreis et al., 2018). According to the current model, an interaction of the short cytoplasmic part of FMG-1B with Intraflagellar Transport (IFT) may occur (Laib et al., 2009; Shih et al., 2013). IFT moves bidirectionally along the flagellar microtubules, the anterograde transport is driven by kinesin 2 and retrograde transport is driven by cytoplasmic dynein-1b (Cole et al., 1998; Huangfu et al., 2003; Kozminski et al., 1995; Lechtreck, 2015; Pedersen and Rosenbaum, 2008; Porter et al., 1999; Rosenbaum and Witman, 2002). Since retrograde motor dynein-1b pauses relative to the adhesion site while FMG-1B tethers to the solid surface through its large extracellular carbohydrate domain (Bloodgood, 2009), the force generated by motor proteins will push the microtubule into the opposite direction, dragging the cell body and the second flagellum behind; the gliding process is initiated (Shih et al., 2013).

Considering that *N*-glycoproteins *per se* are important for adhesion but are constantly lost from the flagellar membrane, gliding is supposed to require an enormous amount of energy, which suggests that flagella mediated adhesion has a somewhat high importance. Furthermore, it opens the question whether the maturation of *N*-glycans (as additional energy expense) in Golgi is important for flagella mediated adhesion, i.e. whether *N*-glycosylation is crucial for adhesion beyond proper glycoprotein folding. Therefore, we compared flagellar membrane motility of different mutant strains impaired in *N*-glycan maturation (characterized in Schulze *et al.* 2018). We found that *N*-glycan maturation indeed impacts the interaction of flagellum and surface in all mutants analyzed.

## Results

### Altered *N*-linked glycans do not change the flagellar localization of FMG-1B

To test whether *N*-glycan maturation in Golgi is important for flagella surface motility in *C. reinhardtii*, two insertional mutants such as IM_*Man1A*_, IM_*XylT1A*_ and their double mutant IM_*Man1A*_xIM_*XylT1A*_ were used. Initially, these mutants had been described in Schulze *et al.* 2018, where *N*-glycan patterns of supernatant proteins were analyzed and compared (Fig. 1A). The insertional mutagenesis giving rise to these mutants was performed in the parental strain CC-4375 (a *ift46* mutant backcrossed with CC124) complemented with IFT-46::YFP referred to as WT-Ins throughout the current study. The first mutant, deficient in xylosyltransferase 1A (IM_*XylT1A*_), produces *N*-glycans devoid of core xylose while simultaneously having a reduced length. Second mutant IM_*Man1A*_, a knock down mutant of mannosidase 1A, is mainly characterized by a lack of 6*O*-methylation of mannose residues while the *N*-glycan length is slightly greater than in WT-Ins. Furthermore, *N*-glycans of IM_*Man1A*_ are slightly reduced in terminal xylose and core fucose. Finally, a double mutant of the above two single mutants (IM_*Man1A*_xIM_*XylT1A*_, obtained by genetic crossing) produces *N*-glycans devoid of 6*O*-methylation but of WT-Ins length and carrying core xylose and fucose residues (Fig. 1A). It is of note that in none of the mutants the flagellar length is altered as compared to WT-Ins (Figure 1 – Supplementary Fig. 1). To confirm that flagellar *N*-glycan patterns of these mutants deviate from WT-Ins, whole cell extracts and isolated flagella were probed with anti-HRP, which is specifically binding to β1,2-xylose and α1,3-fucose attached to the *N*-glycan core (Kaulfürst-Soboll et al., 2011). In line with previous publications, the antibody showed a higher affinity towards *N*-glycoproteins synthesized by IM_*Man1A*_ and IM_*Man1A*_xIM_*XylT1A*_ while it showed a decreased affinity towards probes of IM_*XylT1A*_ (Figure 1 – Supplementary Fig. 2) (Schulze et al., 2018).

**Figure 1.**
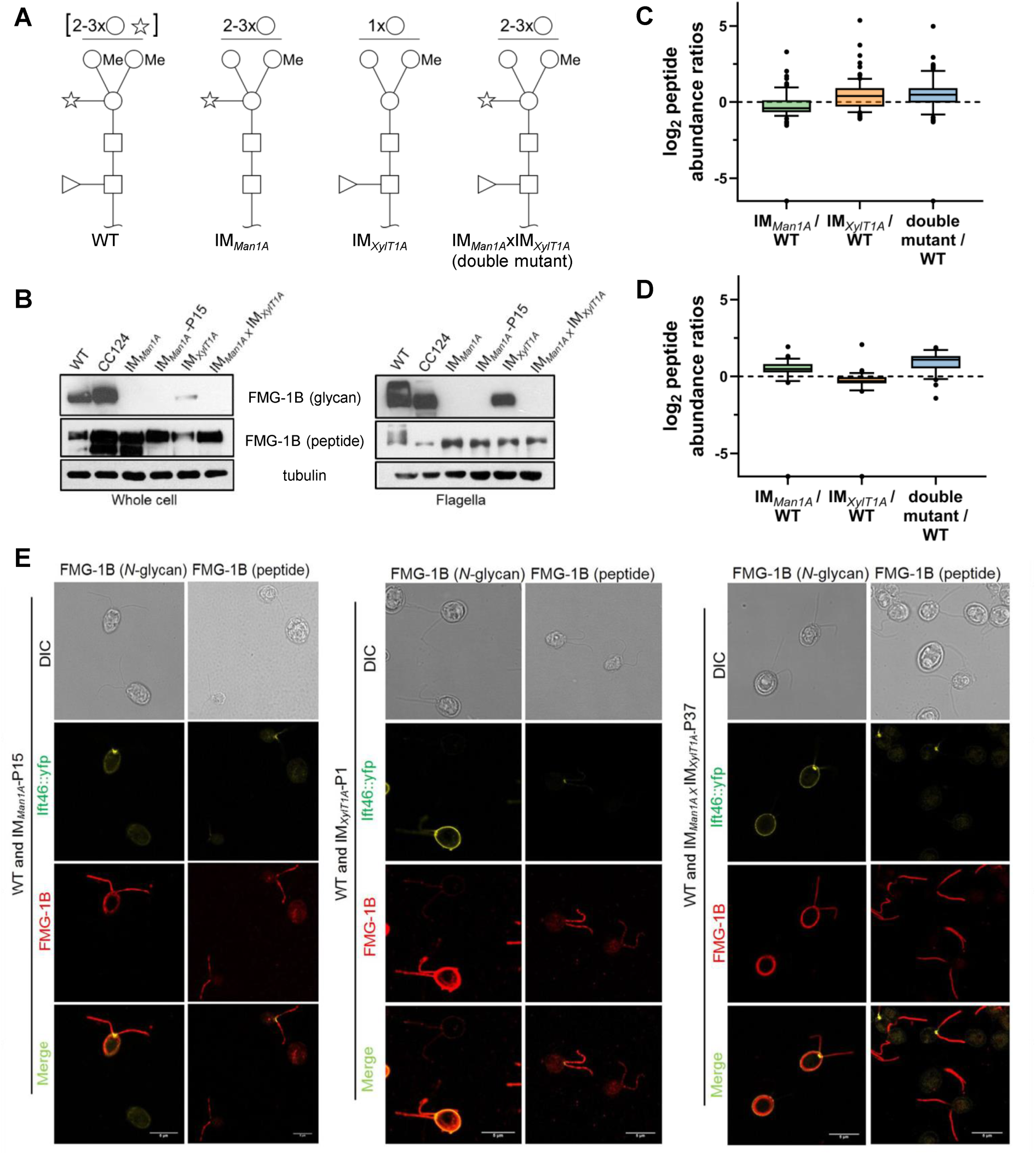
Altered N-glycosylation of FMG-1B does not change its flagellar localization. A, Diagram of N-linked glycan compositions of mutant strains characterized in Schulze *et al.* 2018 and used in the current study. While IM_Man1A_ and the double mutant are mainly characterized by a lower degree of methylation, N-glycans of IM_XylT1A_ are decreased in length and lack the core xylose. All monosaccharides depicted above the horizontal line can be bound to any subjacent residue or to any residue at the same level. Square: N-Acetylglucosamine, Circe: hexose, Triangle: fucose, Star: xylose). B, Probing whole cell and flagella samples with antibodies directed against N-glycans or the protein backbone of FMG-1B.(Bloodgood et al., 1986) C, Log2 peptide abundances of FMG-1B obtained by label free MS analysis. D, Log2 peptide abundances of FAP-113 obtained by label free MS analysis. E, Immuno staining of WT-Ins (IFT46::YFP labelled) cells mixed with mutant cells (IM_Man1A_, IM_XylT1A_ or IM_Man1A_xIM_XylT1A_) with antibodies of FMG-1B.

Since FMG-1B is the major constituent of the flagellar glycoproteome and to date is the only protein proven to be involved in flagellar surface motility, different monoclonal antibodies raised against FMG-1B were employed to analyze FMG1-B localization (Bloodgood et al., 1986; Long et al., 2016). The whole cell or isolated flagella from IM_*Man1A*_, IM_*XylT1A*_ and their double mutant IM_*Man1A*_xIM_*XylT1A*_ were probed separately with the glycan epitope recognizing antibody or the antibody against the peptide epitope of FMG-1B. Both antibodies gave strong signals in WT-Ins and WT-CC-124 (WT-CC-124 is minus gamete used to cross with mutants) (Fig. 1B). On the contrary, the glycan specific antibody showed low affinity towards FMG-1B in cell body and flagella of IM_*Man1A*_, IM_*XylT1A*_ and the double mutant, proving that *N*-glycosylation of FMG-1B is altered in respective strains (Fig. 1B). Additionally, mass spectrometric analyses of intact *N*-glycopeptides derived from the flagella of these mutants confirmed that the change of flagellar *N*-glycan composition is similar to the previously reported composition of *N*-linked glycans among supernatant proteins (Figure 1-Supplementary Fig. 3) (Schulze et al., 2018). Conversely, the antibody specifically binding to the peptide backbone of FMG-1B resulted in signals comparable to WT-Ins for all mutants analyzed. In addition, label free mass spectrometric quantification confirmed that FMG-1B is correctly targeted to the flagella in all mutants analyzed (Fig. 1B and C). The same is true for FAP113, another protein shown to be involved in flagella surface motility (Fig. 1D).

To compare the localization of glycan and protein of FMG-1B in WT-Ins and mutants, IFT46::YFP was removed from the IM strains by crossing the mutant with WT-CC124. The obtained progenies of IM_*Man1A*_, IM_*XylT1A*_ and IM_*Man1A*_xIM_*XylT1A*_ did not express IFT46::YFP (Figure 1 – Supplementary Fig. 4). For imaging, these mutants were mixed with an equal amount of WT-Ins cells and incubated with FMG-1B glycan- or peptide specific antibodies. In the mixed sample of WT-Ins with IM_*Man1A*_, a uniform signal of the glycan specific antibody was found in the flagella and cell wall of WT-Ins. At the same time no glycan signal was observed for the flagella of IM_*Man1A*_ and the signal in the cell wall of IM_*Man1A*_ was very weak. It must be noted, that cross reaction of the *N*-glycan specific antibody with cell wall localized glycosylated proteins has been reported previously.(Bloodgood et al., 1986) On the other hand, the FMG-1B peptide signal concentrated in the flagella of both WT-Ins and IM_*Man1A*_ (Fig. 1E, first column). Similar to IM_*Man1A*_, the glycan signal was not detected in flagella of IM_*XylT1A*_ and the double mutant, while the FMG-1B peptide signal was clearly present in the flagella of these two mutants (Fig. 1E, second and third columns). All these data showed that altered *N*-glycan maturation did not affect the flagellar localization of FMG-1B. Although FMG-1B is the most prominent and best studied flagella membrane glycoprotein, there might be other glycoproteins involved in flagellar adhesion. Nevertheless, no protein was found consistently and significantly changed in abundance in flagella of the IM strains analyzed when compared to WT-Ins (Figure 1 – Supplementary Fig. 5), also indicating that flagellar assembly is not considerably altered in the mutants versus WT.

### Altered *N*-linked glycans attenuate bead attachment and movement to and along the flagellar membrane

In *C. reinhardtii*, polystyrene microspheres adhere to and move bidirectionally along the flagellar surface.(Bloodgood, 1981) To check the effect of altered *N*-glycans on these processes, beads with a diameter of 0.7 μm were added to cell suspensions of IM_*Man1A*_, IM_*XylT1A*_ and the double mutant grown in M1 medium, and the number of beads attached to- or moved along flagella were quantified (Fig. 2A and B). In WT-Ins strains, about 52 % of all flagella had at least one bead attached, whereas the percentage of flagella with beads bound decreased to 46 % in IM_*Man1A*_, 33 % in IM_*XylT1A*_ and 27 % in the double mutant (Fig. 2A). Among the beads attached, 37 % moved along flagella of WT-Ins, 22 % in IM_*Man1A*_, 29 % in IM_*XylT1A*_ and 27 % in the double mutant (Fig. 2B). These data suggested that interaction of flagellar membrane and surface is altered due to altered *N*-glycan composition in these mutants.

**Figure 2.**
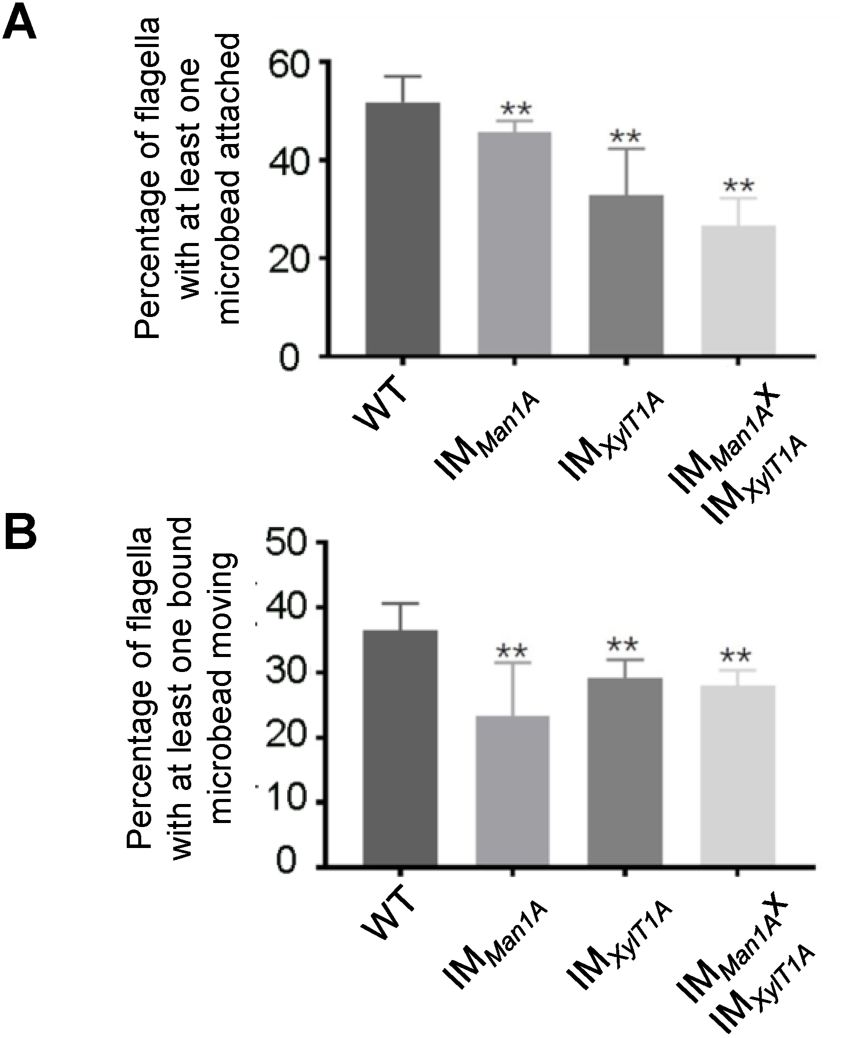
Altered *N*-glycosylation diminishes flagellar polystyrene bead attachment and -transport. A, Percentage of flagella with at least one polystyrene bead bound. Cells were incubated with polystyrene beads (0.7 µm in diameter) and subsequently analyzed by light microscopy. B, Percentage of polystyrene beads transported along the flagellum with at least one polystyrene bead bound. Results present mean of three replicates with 50 cells analyzed per replicate. Error bars show SEM of three replicates. T-test was used for statistical analysis.

### Quantification of the flagella mediated adhesion using atomic force microscopy

Flagella-mediated surface adhesion force was measured via atomic force microscopy (AFM) (Fig. 3A). Cells adhered to a cover slide were attached to an AFM cantilever via physical contact and unspecific binding (Liu et al., 2011). Subsequently, the AFM cantilever was pulled upwards and the force required to pull the cells off was recorded (Fig. 3B). To inhibit whole-cell gliding during AFM pulling, ciliobrevin D was used to inhibit dynein-1b activity and consequently the cell gliding ability (Firestone et al., 2012). Remarkably, the forces necessary to overcome the adhesion of *C. reinhardtii* flagella to the surface were significantly reduced in all IM strains analyzed as compared to WT-Ins (Fig. 3B and C). Especially in the double mutant the average force was reduced from 8 nN in WT-Ins to 1 nN, while the average energy was reduced from 4 to 0.5 J nm^-1^ (Fig. 3C and D). When dynein1-b inhibitor ciliobrevin D was removed, the adhesion force of WT-Ins decreased from 8 nN to 3 nN, while this effect did not occur in the double mutant (Fig. 5A).

**Figure 3.**
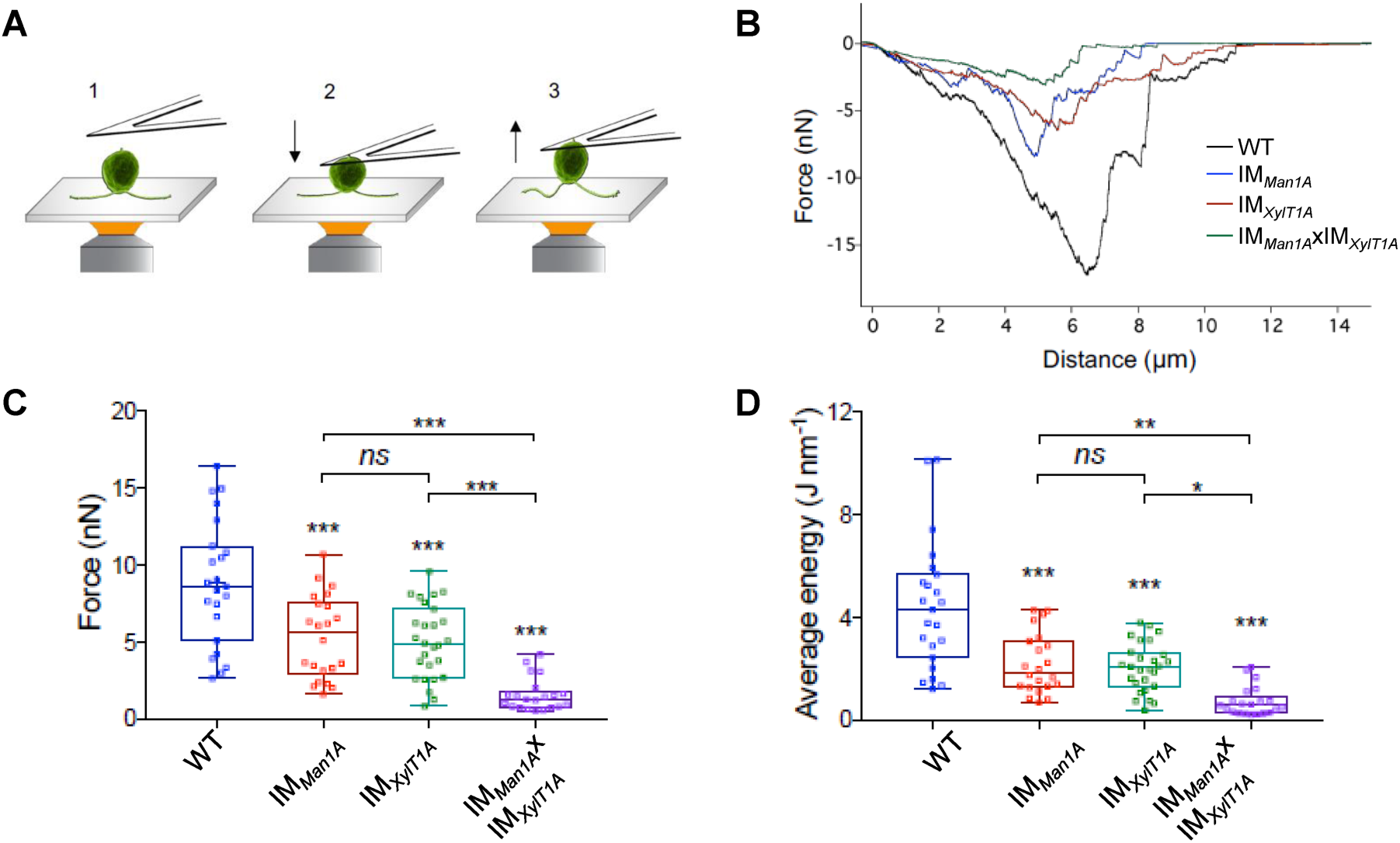
Quantification of the flagella mediated adhesion using atomic force microscopy. A, Diagram of experimental procedures for force measurement: the cell adhered to the surface (1); the cell attached to the AFM cantilever (2); the cell was pulled up from the surface by AFM cantilever (3). Please note, that retrograde IFT, i.e. gliding was inhibited by ciliobrevin D during all measurements presented here. B, Representative force curves acquired for strains including WT, IM_*Man1A*_, IM_XylT1A_ and IM_*Man1A*_xIM_*XylT1A*_. C, Flagella adhesion forces of WT-Ins, IM_*Man1A*_, IM_*XylT1A*_ and IM_*Man1A*_xIM_*XylT1A*_ were generated from force curves (B). D, Average energy of flagellar adhesion of WT-Ins, IM_*Man1A*_, IM_*XylT1A*_ and IM_*Man1A*_xIM_*XylT1A*_. Three biological replicates were performed with minimum 5 cells measured per replicate. *p<0.05, **p<0.01, ***p<0.001. The p-values are obtained from a two-sided, two sample t-test of mean values.

**Figure 4.**
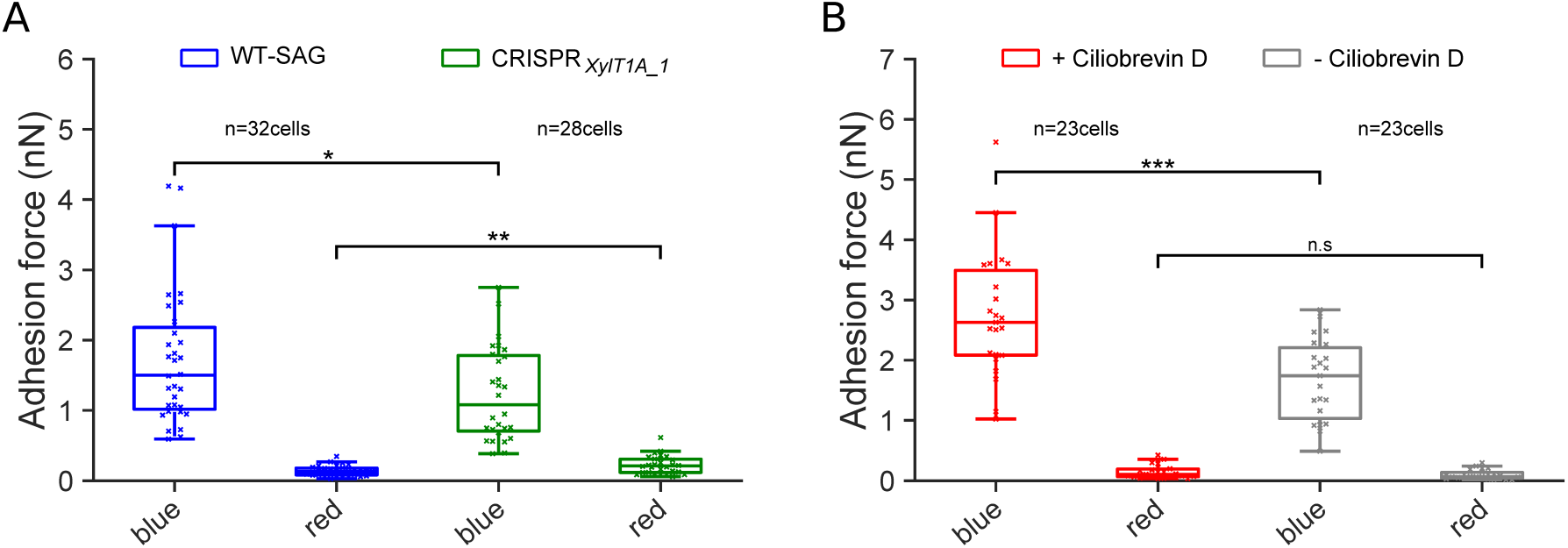
Assessing flagella adhesion forces using micropipette force measurements. A, Flagella mediated adhesion forces acquired for WT-SAG and a xylosyltransferase 1A mutant generated in the genetic background of WT-SAG (CRISPR_*XylT1A_1*_). Micropipette force measurements of the same cells were performed under blue and red light. B, Flagella mediated adhesion forces of WT-SAG in the absence (-) or presence (+) of ciliobrevin D under blue and red light using the same technique. Mean values of 10 measurements per cell are depicted, statistical analysis was performed on mean values: n.s: p>0.05, *p<0.05, **p<0.01, ***p<0.001. The p-values are obtained from Mann–Whitney U test.

**Figure 5.**
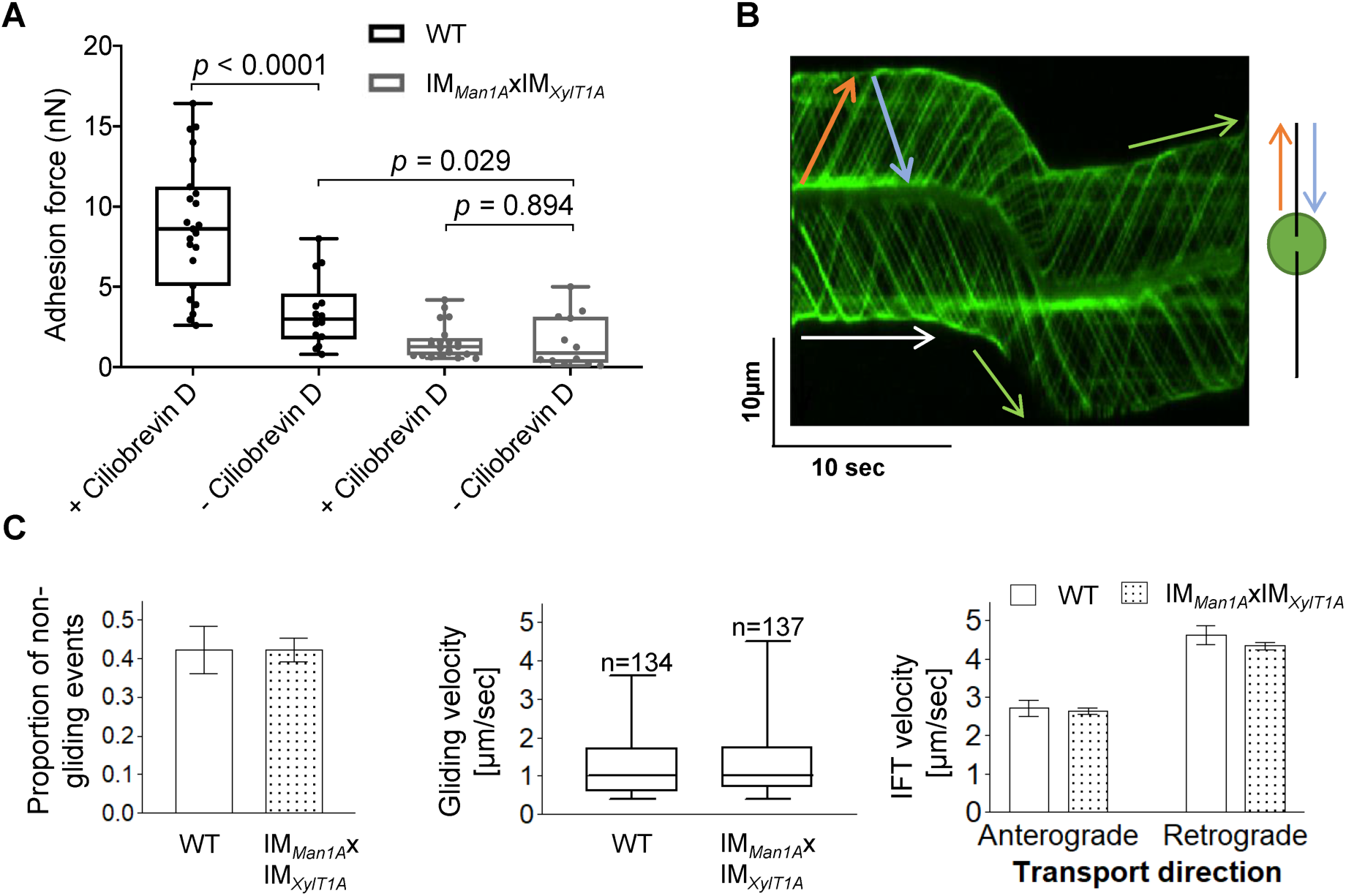
Ciliobrevin D enhances flagella mediated adhesion force in WT. A, Adhesion forces acquired for WT-Ins and the double mutant IM_*Man1*A_xIM_X*ylT1A*_ in the absence or presence of ciliobrevin D, respectively. Three biological replicates were performed with minimum 5 cells measured per replicate. B, Representative kymograph of the movement of IFT46::YFP in flagella of WT-Ins acquired with TIRF microscopy used to calculate the velocity of gliding and IFT. White arrow: non-gliding event (v<0.3µm/s); green arrow: gliding event (v>0.3µm/s); red arrow: anterograde IFT track; blue: retrograde IFT track. C, IFT and gliding are not significantly altered in the double mutant compared to WT-Ins. Proportion of non-gliding events (left), gliding velocity (excluding non-gliding events; middle), IFT velocities in either direction (right). Three biological replicates were performed with 10 cells evaluated per replicate corresponding to 300 IFTs/replicate/strain in case of IFT velocity. Error bars in bar plots represent SD of three replicates. Student-t test was performed comparing mean values of replicates in regard of proportion of non-gliding events (p=0,415) and IFT-velocity (anterograde p= 0,284; retrograde p=0,065). Distribution of gliding velocities was analyzed by use of Mann-Whitney U test (p=0,421), n represents number of gliding events measured.

### Quantification of flagella adhesion using a micropipette force measurement approach

To validate the AFM adhesion force measurements, an independent *in vivo* force measurement approach(Backholm and Bäumchen, 2019; Kreis et al., 2018) was used with another genetic background mutant, a xylosyltransferase 1A (CRISPR_*XylT1A_1*_) mutant, generated in parental wildtype SAG11-32b (WT-SAG) by employing CRISPR/Cas9 (Figure 4 – Supplementary Fig. 1). As a wavelength dependency of flagellar adhesion had been revealed using this approach (Backholm and Bäumchen, 2019), adhesion forces of the same cells were measured under precisely controlled blue- and red-light conditions. Importantly, adhesion forces measured in CRISPR_*XylT1A_1*_ were significantly diminished in comparison to respective WT-SAG, from 1.50 nN to 1.08 nN in blue light (Fig. 4A). Illuminating cells with red light dramatically decreased the adhesion force in both WT-SAG and CRISPR_*XylT1A_1*_. Under red light, mean adhesion forces of individual cells were dispersed broader in CRISPR_*XylT1A_1*_, yet, surface adhesion was enhanced for CRISPR_*XylT1A_1*_ as compared to WT-SAG. In addition, micropipette force measurements under blue light confirm that the presence of ciliobrevin D significantly enhances the adhesiveness of the WT-SAG (Fig. 4B), while no significant effect of ciliobrevin D was observed in adhesion force measurements of the same cells in red light.

### The effect of altered *N*-glycosylation on IFT and gliding

The impact of ciliobrevin D on the adhesion force suggested that IFT might contribute to surface adhesion forces. In addition, it implied that IFT might be hampered via altered *N*-glycosylation. Therefore, IFT velocity and gliding ability of IFT46::YFP expressing WT-Ins and IM_*Man1A*_xIM_*XylT1A*_ were assessed by using total internal reflection fluorescence (TIRF) microscopy. Videos of adhered cells generated by TIRF microscopy were evaluated manually via kymographs in ImageJ (Fig. 5B). Obtained data revealed that neither the proportion of non-gliding events (gliding velocity smaller 0.3 µm*s^-1^), nor gliding speed distribution was significantly diminished when comparing WT-Ins and the double mutant (Fig. 5C). Likewise, anterograde and retrograde IFT velocity was found at WT-Ins levels when comparing WT-Ins and the double mutant (Fig. 5C), implying no significant impact of altered *N*-glycan maturation on IFT.

## Discussion

Our data revealed that the maturation of *N*-glycans has an impact on flagella mediated cell adhesion in *C. reinhardtii.* At the same time, IFT and gliding velocity were not changed due to the altered *N*-glycosylation.

Microbead binding was found diminished in IM strains, implying that the flagellar surface has an altered affinity towards microbeads. In line with this, surface adhesion forces were significantly reduced compared to WT-Ins. The AFM data were confirmed by assessing another XylT1A mutant created via CRISPR/Cas9, using micropipette force measurements. It should be noted that *N*-glycan patterns of IM_*XylT1A*_ and CRISPR_*XylT1A*_ were comparable and thereby strengthen the proposed role of XylT1A as core xylosyltransferase (Lucas et al., 2020; Schulze et al., 2018). Forces measured for both WT strains in the absence of ciliobrevin D were similar in the approaches indicating the comparability of both methods, and confirmed that differential *N*-glycan maturation, i.e. altered *N*-glycan structures attached to mature proteins, lowers the adhesion force of flagella to a surface. These changes in adhesion forces were not accompanied by consistent drastic changes in the flagellar proteomes. For example, FMG-1B and FAP113, to date the only two known proteins involved in surface adhesion, were found in comparable amounts in WT-Ins and mutants (Fig. 1 and Figure 4 – Supplementary Fig. 2) (Bloodgood et al., 2019; Kamiya et al., 2018). Of note, also FMG-1A is localized in flagella and its amount was unaltered between WT and mutants in vegetative cells (Figure 1 – Supplementary Fig. 3). Interestingly, gliding of mutant strains on solid surface was not affected. The current model for flagella mediated cell adhesion and subsequent gliding proposes that the extracellular part of certain glycoproteins such as FMG-1B adheres to the surface, cytoplasmic moieties of these proteins are bound to ongoing retrograde IFT directly or indirectly upon calcium- and light dependent stimulus which is followed by an onset of gliding (Kreis et al., 2018; Shih et al., 2013). Assuming that altered *N*-glycan maturation does not impact initial protein folding in the ER (as those steps are spatially and temporally separated), our data revealed that changed *N*-glycosylation, did not alter targeting of respective glycoproteins to flagella nor the velocity of IFT. Thus, changes in surface adhesion are likely linked to *N*-glycoprotein epitopes and their direct interaction with the solid or semisolid surface.

Notably, ciliobrevin D enhanced adhesion forces measured for WT-Ins but not for the double mutant. The difference may result from the activity of cytoplasmic dynein-1b, retrograde motor of IFT. As IFT and gliding were not changed between double mutant and WT-Ins (Fig. 5), the differential ciliobrevin D dependent adhesion forces imply that the ciliobrevin D mediated enhancement of binding forces is not entirely due to IFT inhibition but more complex. Although it is well accepted that ciliobrevin D specifically inhibits dynein-1b function, abolishes retrograde IFT and is therefore widely used in cilia/flagella research, strikingly little is known about its mode of action.(Firestone et al., 2012) When first described, it was suggested that it would impair ATPase activity and not much knowledge was gained ever since (Roossien et al., 2015). Clearly, more work is required to understand the mechanics of the ciliobrevin D-induced enhancement of adhesion force (Fig.4 and 5).

The fact that the altered *N*-glycosylation diminished the force of cells to adhere to surfaces but did not affect IFT, strongly suggests that adhesion to surfaces and IFT are not necessarily coupled. As discussed below, adhesion probably evolved independently of the necessity to enable cell gliding.

As described before, the adhesion forces for both strains were significantly stronger under blue than under red light (Fig. 4A). Under blue light, the adhesion force for CRISPR_*XylT1A*_ was significantly weaker compared to WT-SAG, consistent with what was observed via the AFM force measurements (Fig. 3). Momentarily, there are no mechanistic insights regarding the light dependence of *C. reinhardtii* adhesion and also the functional blue-light photoreceptor remains to be identified. Yet, the red-light suppression of adhesion is diminished in CRISPR_*XylT1A*_, indicating that *N*-glycosylation might also impact flagellar driven processes other than adhesion.

In summary, taking advantage of single-cell adhesion force measurements, our data revealed that cell adhesion was significantly impaired in *C. reinhardtii N*-glycosylation mutant strains. Our data further suggested that flagellar assembly, IFT and FMG-1B transport into flagella were not affected by altered *N*-glycosylation. Thus, we conclude that proper *N*-glycosylation of flagellar proteins is crucial for adhering *C. reinhardtii* cells onto surfaces. We further suggest that the remaining adhesion force, although diminished in *N*-glycan mutants, is still sufficient for gliding. This implies that the evolution of adhesion was not governed by the capability of gliding but adhesion itself. Given the response of flagellar adhesion blue light, it could potentially link adhesion to photo-protection which is also blue-light mediated, as blue light mediated adhesion might result in photoprotection via cell shading (Kreis et al., 2018; Petroutsos et al., 2016).

## Material and Methods

### Culture growth

Cells were grown photoheterotrophically in tris-acetate-phosphate (TAP) medium under constant illumination at 50 µmol photons*s^-1^*cm^-2^ unless stated otherwise.

### Measurement of flagellar length and flagellated cells

The cells were fixed with 0.5% Lugol’s solution for 1h at room temperature. Flagellar length measurements were performed using a phase microscope (Nikon Eclipse Ti) equipped with an electron multiplying charged-coupled device. For each sample, at least 50 flagella were measured. For the measurement of flagellated cells, at least 100 cells were counted for each strain in biological triplicates.

### Generation and analysis of a CRISPR/Cas9 mutant strain

Mutagenesis was performed on the WT strain SAG11-32b following the protocol described in Greiner *et al.* 2017 employing the transformation of a pre-built Cas9:guideRNA complex (Cas9 target sequence in the XylT1A gene: ACGAACACCCCAACACCAAT) simultaneously with a plasmid encoding for a paromomycin resistance via electroporation. Following selection with paromomycin, putative mutants were screened by PCR using the primer pairs short_fw: TACAAAGAACGGGACGCAGG, short_rev: CATTGAAGCTCATCCAGACAC and long_fw: AAGGGTCACGGCACGGTATG, long_rev: CCTGAAGCACCCATGATGCACG. Genomic *XylT1A* regions of candidate strains showing not-WT like band patterns were sequenced. In total, two mutant strains differing in the DNA inserted following the Cas9 cutting site were identified (CRISPR_*XylT1A_1*_ and CRISPR_*XylT1A_2*_). Next, XylT1A protein levels were quantified by parallel reaction monitoring (PRM) and supernatant *N*-glycan compositions were assessed by IS-CID mass spectrometry. Additionally, flagella were isolated, separated by SDS-PAGE and, after transferred to a nitrocellulose membrane, probed with the protein backbone FMG-1B specific antibody.

### Flagella isolation

Flagella isolation from cultures in the mid-log growth phase was performed as described elsewhere by the pH shock method (Witman et al., 1972). Pellets containing flagella samples were stored at −80°C until further use for immunoblotting or sample preparation for mass spectrometric measurements.

### Immunoblotting

Frozen, dry flagella and whole cell samples were resuspended in lysis buffer (10mM Tris/HCl, pH=7.4, 2 % SDS, 1 mM Benzamidine and 1 mM PMSF) and subjected to sonication for 10 min. After pelleting the not-soluble cell debris, the protein concentration was determined using the bicinchoninic acid assay (BCA Protein Assay Kit by Thermo Scientific Pierce). Volumes corresponding to 30 µg of protein were separated by SDS-PAGE, transferred to nitrocellulose membranes and incubated with antibodies as indicated.

### Sample preparation for mass spectrometric measurements

Frozen, dry flagella and whole cell samples were treated as described in *Immunoblotting*. Volumes corresponding to 60 µg of protein were tryptically digested and desalted as described elsewhere (Rappsilber et al., 2007).

### Mass spectrometry measurements

Tryptic peptides were reconstituted in 2 % (v/v) acetonitrile/0.1 % (v/v) formic acid in ultrapure water and separated with an Ultimate 3000 RSLCnano System (Thermo Scientific). Subsequently, the sample was loaded on a trap column (C18 PepMap 100, 300 µm x 5 mm, 5 mm particle size, 100 Å pore size; Thermo Scientific) and desalted for 5 min using 0.05 % (v/v) TFA/2 % (v/v) acetonitrile in ultrapure water with a flow rate of 10 µL*min^-1^. Following, peptides were separated on a separation column (Acclaim PepMap100 C18, 75 mm i.D., 2 mm particle size, 100 Å pore size; Thermo Scientific) with a length of 50 cm. General mass spectrometric (MS) parameters are listed in Figure 1 – Supplementary Table 1.

For quantification of glycosyltransferases, PRM (including a target list) was employed on whole cell samples and respective spectra were analyzed with the Skyline software (Pino et al., 2017). For quantification of flagellar proteins, flagella samples were measured in biological quadruplicates in standard, not targeted, data dependent measurements. Following, peptide wise protein abundance ratios (IM/WT) were calculated with ProteomeDiscoverer™ (normalizing on a set of not membrane standing flagellar proteins) and filtered for proteins identified in at least 11 samples, for proteins having Abundance Ratio Adj. P-Value < 0.05 for at least one ratio and for proteins appearing in the flagellar proteome ChlamyFPv5.(Pazour et al., 2005)

In order to assign glycopeptides, samples were measured employing In-Source collision induced dissociation (IS-CID) as described previously followed by analysis of data with Ursgal and SugarPy (Kremer et al., 2016; Oltmanns et al., 2020; Schulze et al., 2020).

### Microbead measurements

Microbead binding- and transport assays were performed analogous to previous descriptions.(Bloodgood et al., 2019) Monodisperse polystyrene microspheres (0.7 μm diameter) were purchased from Polysciences, Inc. Beads were washed with deionized water for three times and resuspended in NFHSM to make a store solution, which was used at 1:10 dilution in adhesion and motility detecting experiment.

To quantify the ability of bead binding, beads were added to 500 μL of cells at a density of 2 x 10^7^ cells*mL^-1^. After 5min, cells were observed with a light microscope (Olympus, U-HGLGPS, 100X oil objective). A flagellum was scored as “+ bead” if beads adhered to it. The percentage of flagellar binding beads was calculated as: Percentage of flagellar binding beads = the number of “+ bead”/ (total number flagella scored) x 100%.

To obtain a kinetic measure of surface motility, cells were mixed with beads as above for 5 min and randomly observed under the light microscope. Each bead adhered to a flagellum was monitored for about 30 seconds. If beads moved along the flagella, we marked it as “Moved bead” or it was “Adhered bead”. The surface motility was calculated as: Percentage of moved beads along with flagella= “Move bead” x/ (“Moved bead” + “Adhered bead”) 100%.

### AFM measurements

*C. reinhardtii* strains, grown in M1 medium under constant white illumination were grown for 65 h, were allowed to adhere to a glass slide (immersed in ethanol for overnight, subsequently rinsed with MQ water) in fresh M1 medium for 15 min. Following, cells were incubated in the presence of ciliobrevin D for 1 h (500 µL M1 supplemented with 200 µM ciliobrevin D). For AFM measurements, only adhered cells in gliding conformation having approximately similar appearance were analyzed. The MLCT-O10 AFM probe (Spring Const.: 0.03 N m^-1^, length: 215 µm, width: 20 µm, resonant freq.: 15 kHz, Bruker) was soaked in acetone for 5 min, then subjected to UV illumination (distance to lamp: 3-5 mm) for 15 min. Then, the probe was immersed in 0.01 % poly-l-lysine for 1 h and afterwards rinsed with MQ water. Following, the probe was immersed with 2 % glutaraldehyde for 1 h and rinsed with MQ water before use. The AFM measurement was performed in Force Spectroscopy Mode in liquid at room temperature using a NanoWizard 3 AFM (JPK) equipped with a CellHesion stage (Z range: 100 µm) NanoWizard 3 head. The tip spring constant was routinely calibrated. The AFM tip, modified as described, was lowered onto the cell surface at a rate of 10 μm s^-1^ with a z scale of 25 μm. After contact, the applied force was maintained at 3 nN for 15 s. Then, the cell-attached probe was upraised at a rate of 1 μm s^-1^. Force curves were processed with JPK SPM Data Processing (JPK). The forces and energy were determined as described in Figure 3 – Supplemental Figure 1 (Liu et al., 2011). Three biological replicates were performed with minimum 5 cells measured per replicate.

### Micropipette force measurements

Culture growth and micropipette force measurements were performed following established recipes.(Kreis et al., 2019, 2018) In brief, *C. reinhardtii* strains WT-SAG and CRISPR_*XylT1A_1*_ were grown axenically in tris-acetate-phosphate (TAP) medium (Thermo Fisher Scientific) in a Memmert IPP 100Plus incubator on a 12h day / 12h night cycle. The experimental approach is based on the use of a homemade micropipette force sensor, which allows for grasping a living cell by suction (Backholm and Bäumchen, 2019). The adhesion force is obtained by bringing the flagella into contact with a cleaned piece of a silicon wafer (Si-Mat, unilateral polished), and by measuring the micropipette deflection during iterative approach and retraction of the substrate. The illumination wavelength for blue and red light was 470 nm and 671 nm, respectively, and realized by using narrow band pass interference filters (FWHM: 10 nm). During the adhesion force measurements, the cells were illuminated with a constant photon flux of 10^19^ photons m^-2^ s^-1^ for both light conditions. For each cell, 10 adhesion force measurements were performed for each wavelength, whereby the order of red and blue light was varied randomly after 5 consecutive measurements. For testing the influence of ciliobrevin D, a 200 μM solution of ciliobrevin D (Merck, Darmstadt, Germany) was prepared in a 9:1 water-dimethylsulfoxid (DMSO, Sigma-Aldrich, Germany, Purity: 99.9%) mixture. *C. reinhardtii* cells were incubated in the ciliobrevin D solution for 30 minutes. Since a high DMSO concentration in the TAP medium causes death of the cells, the volume fraction of DMSO was kept at a level of about 3.6% during the incubation and experiments. After the incubation, the culture was centrifuged at 100 *g* for ten minutes to concentrate dead cells at the bottom of the incubation flask. The culture was then rested in the incubator before the experiments for at least 30 minutes. At last, the upper part of the culture in the flask was used to fill the liquid cell to perform the experiments. In parallel, a second sample of *C. reinhardtii* cells was incubated with the same fraction of DMSO (without ciliobrevin D) and followed by the same experimental procedure to serve as a control group.

### TIRF imaging

Total internal reflection microscopy (TIRF) was applied to assess intraflagellar transport and gliding behaviour of *C. reinhardtii* strains expressing YFP-coupled IFT46. Therefore, cell densities were adjusted to 1*10^5^ cells*ml^-1^. Samples were loaded to a glass bottom microscopy chamber (µ-Slide 8 Well Glass Bottom) and refreshed every 20 min while imaging. TIRF microscopy was performed at room temperature with a Nikon Eclipse Ti and a 100x objective. IFT46::YFP was excited at 488 nm and fluorescence was recorded with an iXon Ultra EMCCD camera (Andor). For analysis, images were captured with NIS-Elements software over 30 s at 10 fps and a pixel size of 0.158 µm*pixel^-1^. Images were evaluated by use of Fiji via manual evaluation of kymographs. Nett IFT velocities during gliding were calculated by subtracting corresponding gliding velocities. Three biological replicates were performed with 10 cells in gliding configuration analysed per replicate.

### Confocal Imaging

Cells were incubated with primary antibodies (FMG-1B #8 and #61, available at dshb.com), subsequently incubated with a fluorescently labeled secondary antibody and analyzed by confocal microscopy as described previously (Lv et al., 2017). In brief, cells were plated on 1 % poly(ethyleneimine) coated cover glass, decolorized and fixed in methanol at −20 °C for 20 min, permeated cells in PBS buffer for 1 h, and then blocked in 5 % BSA (Biosharp), 10 % normal goat serum (Dingguo) and 1 % fish gelatin (Sigma) in PBS. Incubated the samples with primary antibodies overnight, washed them, and incubated secondary antibody, washed the samples and mounted them on slides with nail polish.

## Acknowledgements

The work in the laboratory of K.H. was supported by the National Nature Science Foundation of China (Grant 31671399 to Huang K). M.H. acknowledges support from the German Science Foundation (DFG, HI739/12-1). L.N.L. acknowledges funding support from the Royal Society (UF120411, URF\R\180030, RGF\EA\181061 and RGF\EA\180233) and the Biotechnology and Biological Sciences Research Council (BB/R003890/1, BB/M012441/1). China Postdoctoral Science Foundation Funded Project (Project No.: 2019M662335). A.O. and J.B. thank S. Schulze for providing SugarPy. A.G., M.K. and O.B. thank M. Lorenz and the Göttingen Algae Culture Collection (SAG) for providing the WT-SAG strain and R. Catalan for technical assistance.

## Author contributions

N.X and L.H. performed light microscope imaging. A.O., J.B. and M.S. performed mass spectrometry assisted data analysis. N.X., A.O. and J.B. performed immuno blotting experiments. A.O. and S.K. performed CRISPR-Cas mutagenesis with help of P.H.. L.Z and L.L performed AFM measurements and corresponding data interpretation. A.G., M.K. and O.B. performed and analyzed micropipette force measurements. A.O., N.X., L.H., K.H. and M.H. were involved in data interpretation. A.O. wrote the manuscript with help of N.X., L.H., M.H. and K.H.

## Materials and Correspondence

The mass spectrometry proteomics data have been deposited to the ProteomeXchange Consortium (http://proteomecentral.proteomexchange.org) via the PRIDE partner repository with the dataset identifier PXD018353 and will be publically available upon acceptance of the manuscript (Perez-Riverol et al., 2019). During peer review, the dataset can be entered via the account and the password OLy6xQJY. For further requests, please contact K. Huang or M. Hippler.

## Supplementary Material corresponding to the publication

**Figure 1 – Supplementary Table 1.** MS parameters. Relevant parameters used to acquire IS-CID and not fragmented TopN MS spectra as well as PRM data.

|  | TopN without IS-CID | In-Source CID HCD | Parallel reaction monitoring |  |
| --- | --- | --- | --- | --- |
| **Eluent compositions** | Peptide trapping: 0.05% trifluoroacetic acid (TFA) in ultrapure water (A1), 0.05% TFA in 80% acetonitrile (B1)<br>Peptide separation: 0.1% formic acid (FA) in ultrapure water (A2), 0.1% FA in 80% acetonitrile (B2) |  |  | **LC parameters** |
| **Trap Column** | C18 PepMap 100, 300 μM x 5 mm, 5 μm particle size, 100 Å pore size; Thermo Scientific |  |  |  |
| **Peptide trapping (eluent A1+B1)** | 2.5% B1 at 5 μl/min for 5 min |  | 2.5% B1 at 10 μl/min for 3 min |  |
| **Flow rate** | 300 nL/min |  | 250 nL/min |  |
| **Separation Column** | Acclaim PepMap C18, 75 μm x 50 cm, 2 μm particle size, 100 Å pore size; Thermo Scientific |  |  |  |
| **Gradient for peptide separation (eluent A2+B2)** | 2.5% B2 over 5 min,<br>2.5-45% B2 over 40 min,<br>45%-99 % B2 over 5 min<br>99% B2 for 20 min<br>99%-2.5% over 5 min<br>2.5% for 30 min |  | 2.5% B2 over 5 min,<br>2.5-35% B2 over 105 min,<br>35%-99 % B2 over 5 min<br>99% B2 for 20 min<br>99%-2.5% over 5 min<br>2.5% for 40 min |  |
| **In-source CID** | off | 80 eV | off | **MS1 settings** |
| **Use lock masses** | off |  | on (m/z 445.12003) |  |
| **Resolution at m/z 200 (FWHM)** | 70,000 |  |  |  |
| **Chromatographic peak width** | 15 s |  |  |  |
| **AGC target** | 3e6 |  |  |  |
| **Maximum injection time** | 100 ms |  | 50 ms |  |
| **Scan range** | 600-3000 m/z |  | 350-1600 m/z |  |
| **Mass tags** | off | on | off |  |
| **TopN** | 12 |  | n/a | **MS2 settings** |
| **Resolution at *m/z* 200 (FWHM)** | 17,500 |  | 35,000 |  |
| **Isolation window** | 2 m/z |  | 2 m/z (offset 0.5 m/z) |  |
| **AGC target** | 1e5 |  |  |  |
| **Maximum injection time** | 120 ms |  |  |  |
| **Normalized collision energy (NCE)** | 30 |  | 27 |  |
| **Minimum AGC target** | 1.25e3 |  | n/a |  |
| **Intensity threshold** | 1e4 |  | n/a |  |
| **Charge exclusion** | unassigned, >5 |  | n/a |  |
| **Dynamic exclusion** | 15 s |  | n/a |  |

**Figure 1 – Supplementary Figure 1.**
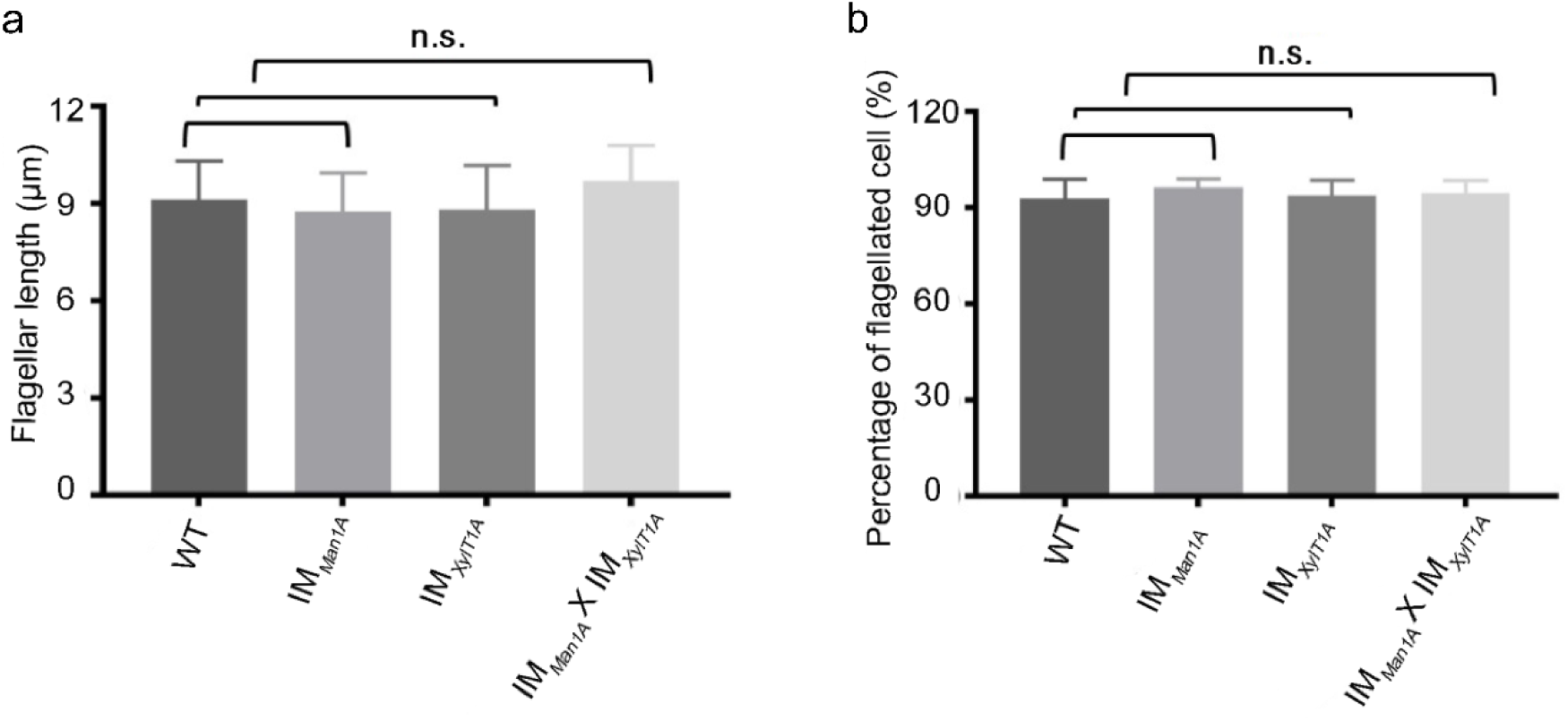
Flagellar length is not altered in *N*-glycosylation mutants. a, Measurement of the flagellar length (in µm) among WT-Ins and three mutants. 50 flagella were measured in each experiment and this experiment has three biological repeats. Error bar: mean ± SD. p>0.05. b, Percentage of flagellated cell in WT-Ins and IM strains. The result of three biological replicates is showns. Error bar: mean ± SD. p>0.05.

**Figure 1 – Supplementary Figure 2.**
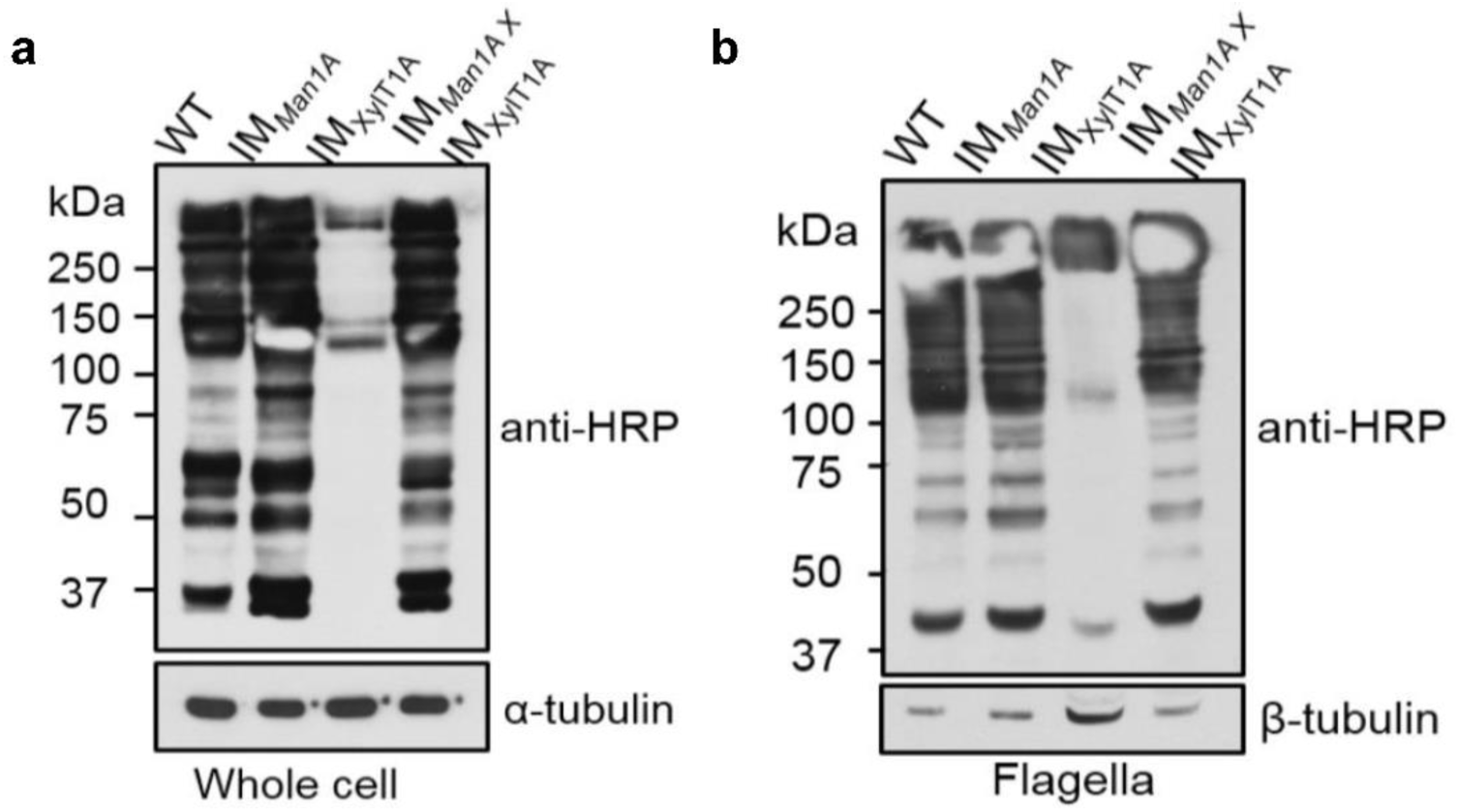
*N*-Glycan structures are altered in mutants as compared to WT-Ins. Whole cell proteins (a) and isolated flagella (b) of WT-Ins and mutants were separated on a 7% SDS-PAGE, transferred to nitrocellulose and probed with anti-HRP, which is specifically binding to β1,2-xylose and α1,3-fucose attached to the *N*-glycan core. α-tubulin, as loading controls.

**Figure 1 – Supplementary Figure 3.**
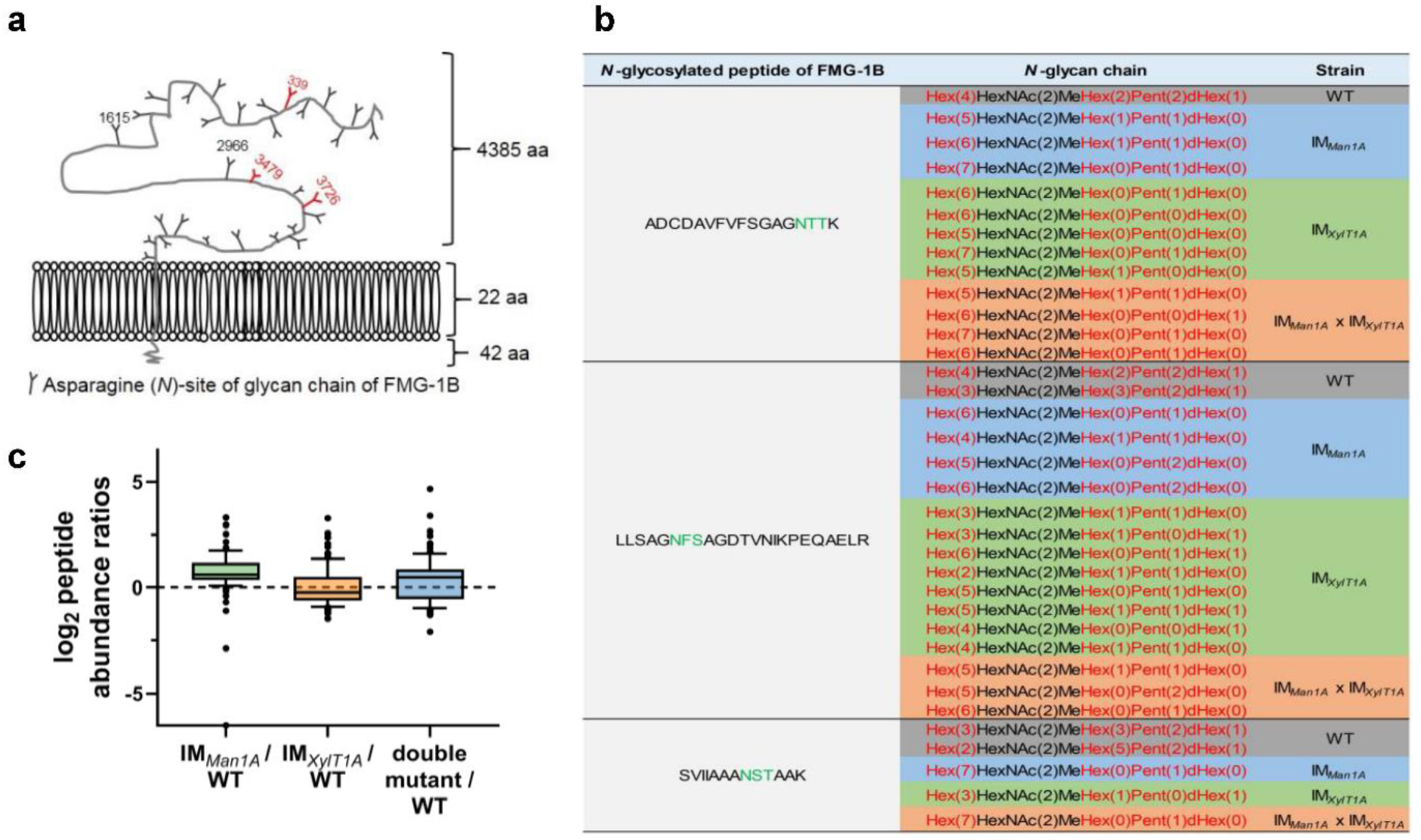
Change of *N*-glycan pattern of FMG-1B in IM strains as compared to WT-Ins. a, Diagram of the topology of FMG-1B, the major component of the glycocalyx in *C. reinhardtii*. The identified *N*-linked glycosylation sites are marked. b, Comparison of FMG-1B proteotypic *N*-glycopeptides identified by IS-CID/SugarPy in WT-Ins, IM_*Man1A*_, IM_*XylT1A*_ and IM_*Man1A*_xIM_*XylT1A*_. c, The relative peptide abundances of FMG-1A in IM_*Man1A*_, IM_*XylT1A*_ and IM_*Man1A*_xIM_*XylT1A*_ strains compared it in WT-Ins obtained by label free MS analysis.

**Figure 1 – Supplementary Figure 4.**
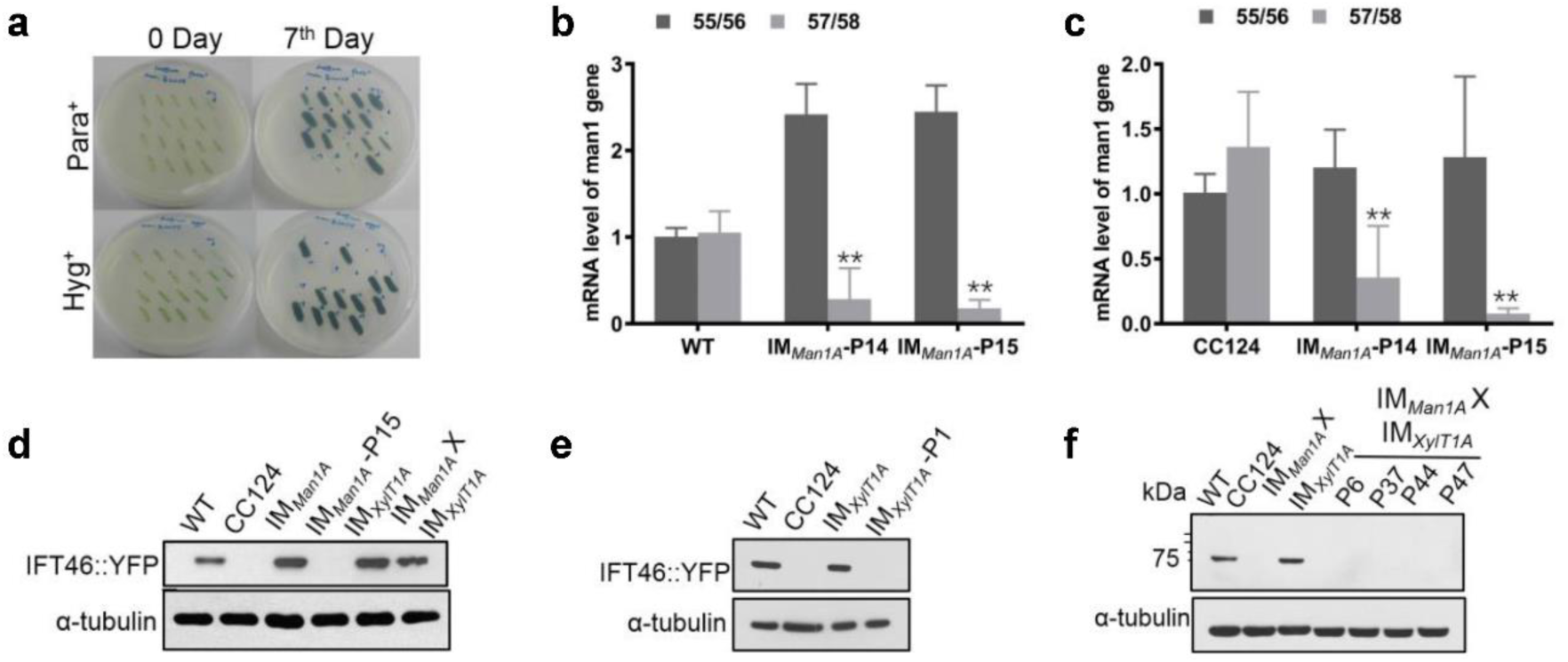
Genetic crossing of original IM strains (mt+) with CC124 (mt-) to obtain mutants lacking IFT46::YFP. a, Screening the progenies of IM_*Man1A*_ mutants with WT-CC-124 to obtain the mutant without IFT46::YFP background, which grew in TAP plate with Paromomycin added and died in TAP plate with Hygromycin added. b, Determination of mRNA levels of *MAN1A* gene in progeny clones P14, P15 and WT-Ins. IM_*Man1A*_ *-*P14 still can live in TAP plate with Paromomycin or Hygromycin. IM_*Man1A*_ -P15 grew in TAP plate with Paromomycin and died in TAP plate added Hygromycin, which is the strain IFT46::YFP had been removed. Error bar: mean ± SD. Asterisks indicate statistically significant differences (*p* < 0.01). c, Comparison of mRNA level of *MAN1A* gene in progeny clones with CC-124. Error bar: mean+/− SD. Asterisks indicate statistically significant differences (*p* < 0.01). d, Determination of the expression of IFT46::YFP in whole cell of WT-Ins and IM_*Man1A*_ -P15 mutant. IM_*Man1A*_ -P15 is the offspring generated by crossing IM_*Man1A*_ -P15 mutant (mt+) with CC124 (mt-). α-tubulin, as loading control. e, Determination of the expression of IFT46::YFP in whole cell of WT-Ins and IM_*XylT1A*_ -P1 mutant. α-tubulin, as loading control. f, Determination of the expression of IFT46::YFP in progenies from the crossing CC124 with IM_*Man1A*_ X IM_*XylT1A*_. α-tubulin, as loading control.

**Figure 1 – Supplementary Figure 5.**
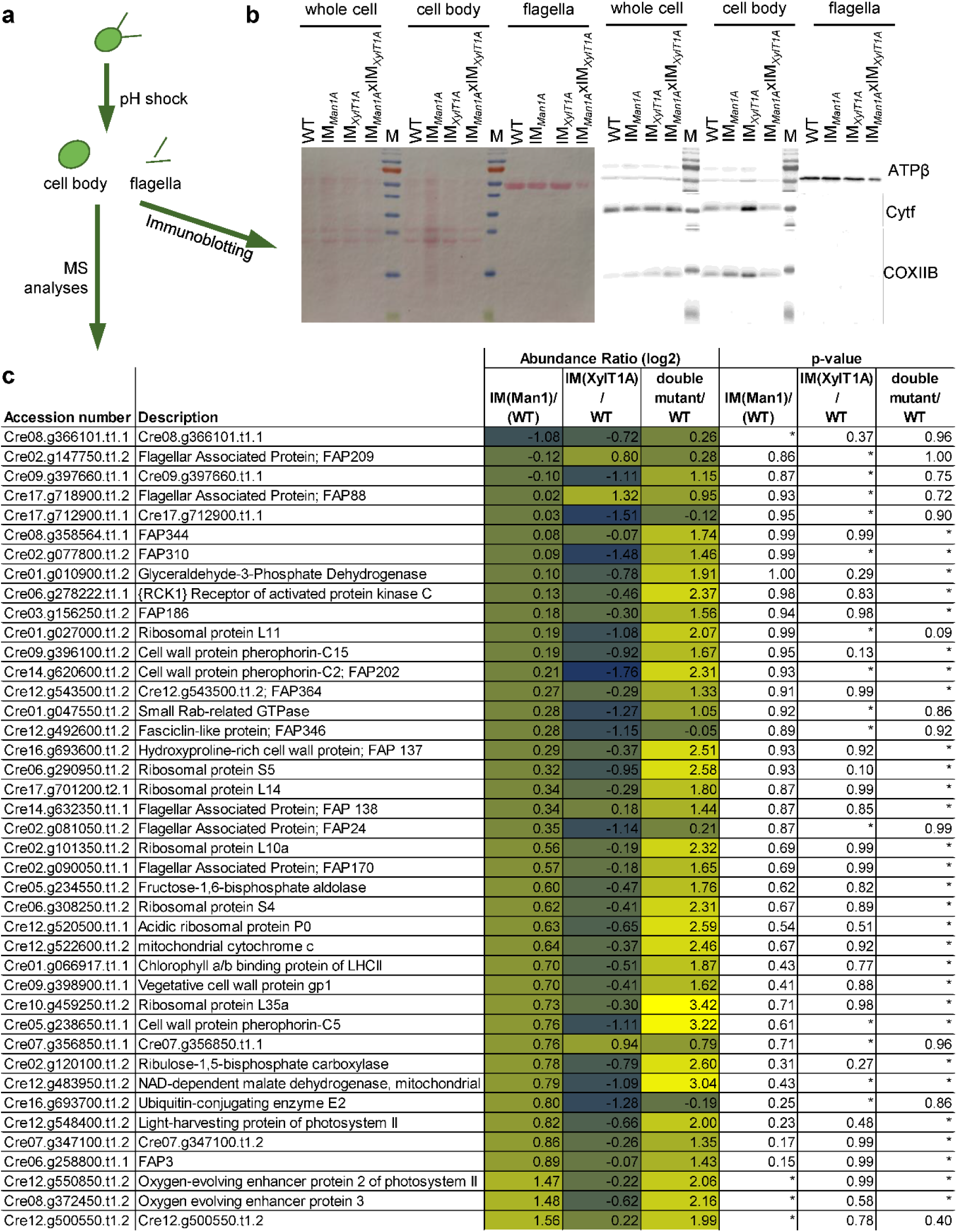
Quantitative mass spectrometry of isolated flagellar from WT-Ins and *N*-glycosylation mutants. a, Experimental procedures of flagella isolation and following analyses. b, Immunoblot of isolated flagella probed with antibodies against chloroplast marker protein (Cytf, cytochrome f) and mitochondrium marker protein (COXIIB). The absence of respective marker proteins in the flagellar fraction prove the purity of flagellar samples while ponceau staining of the same membrane indicates that similar amounts of protein were loaded. c, Proteins found significantly differential in abundance in IM strains compared to WT-Ins. Entries of proteins are given with corresponding phytozome ID protein description.

**Figure 3 – Supplementary Figure 1.**
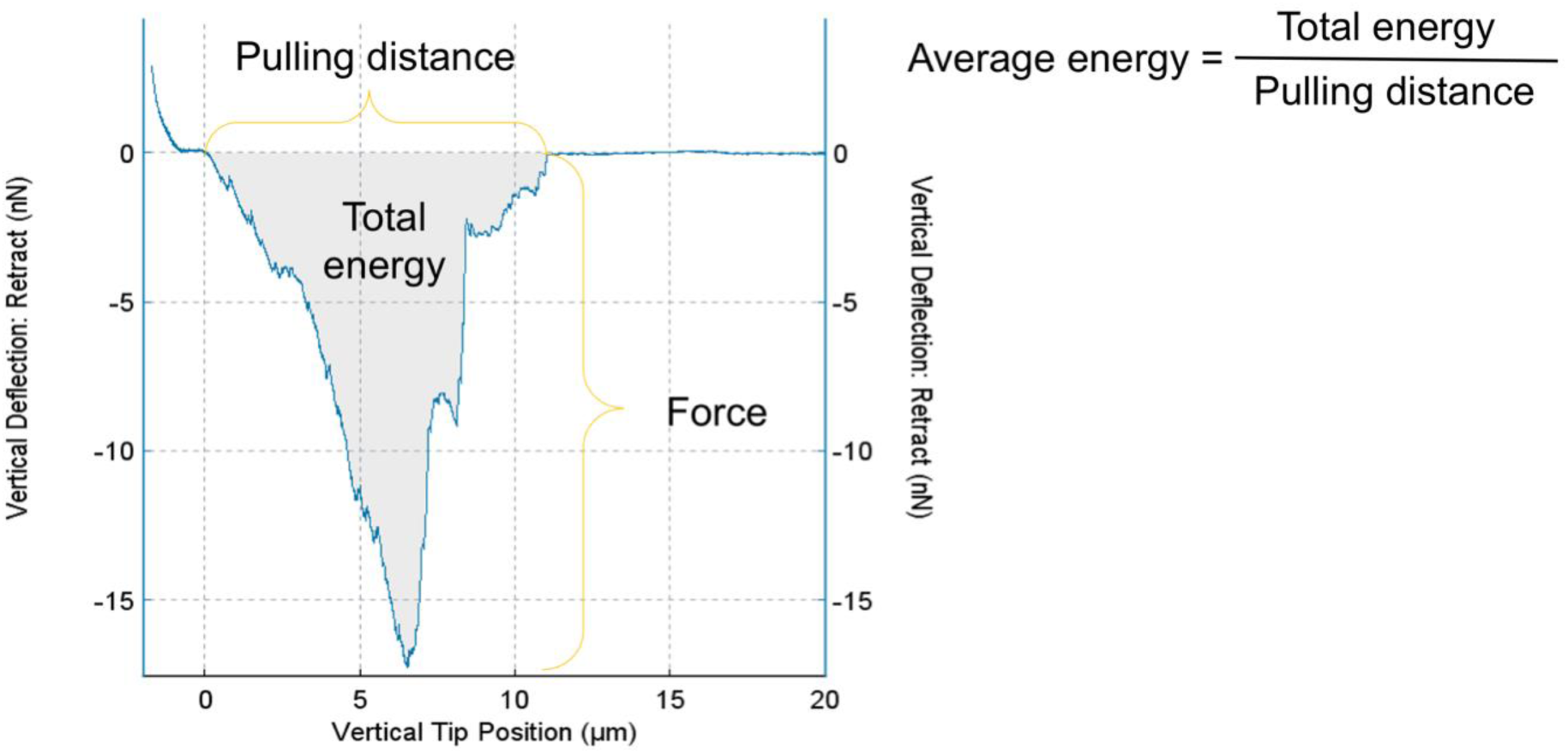
Analysis of the force and energy required to overcome the adhesion of *C. reinhardtii* flagella to the surface from AFM force curves. The curve shows the relation of pulling force and pulling distance during AFM probe retracting. The lowest point of the curve represents the maximal adhesion force of flagella which was used in Figure 3C. The area of shaded region represents the total energy calculated by the JPK SPM Data Processing software for pulling up the cell. The average energy is calculated via dividing total energy by pulling distance as shown in the figure.

**Figure 4 – Supplementary Figure 1.**
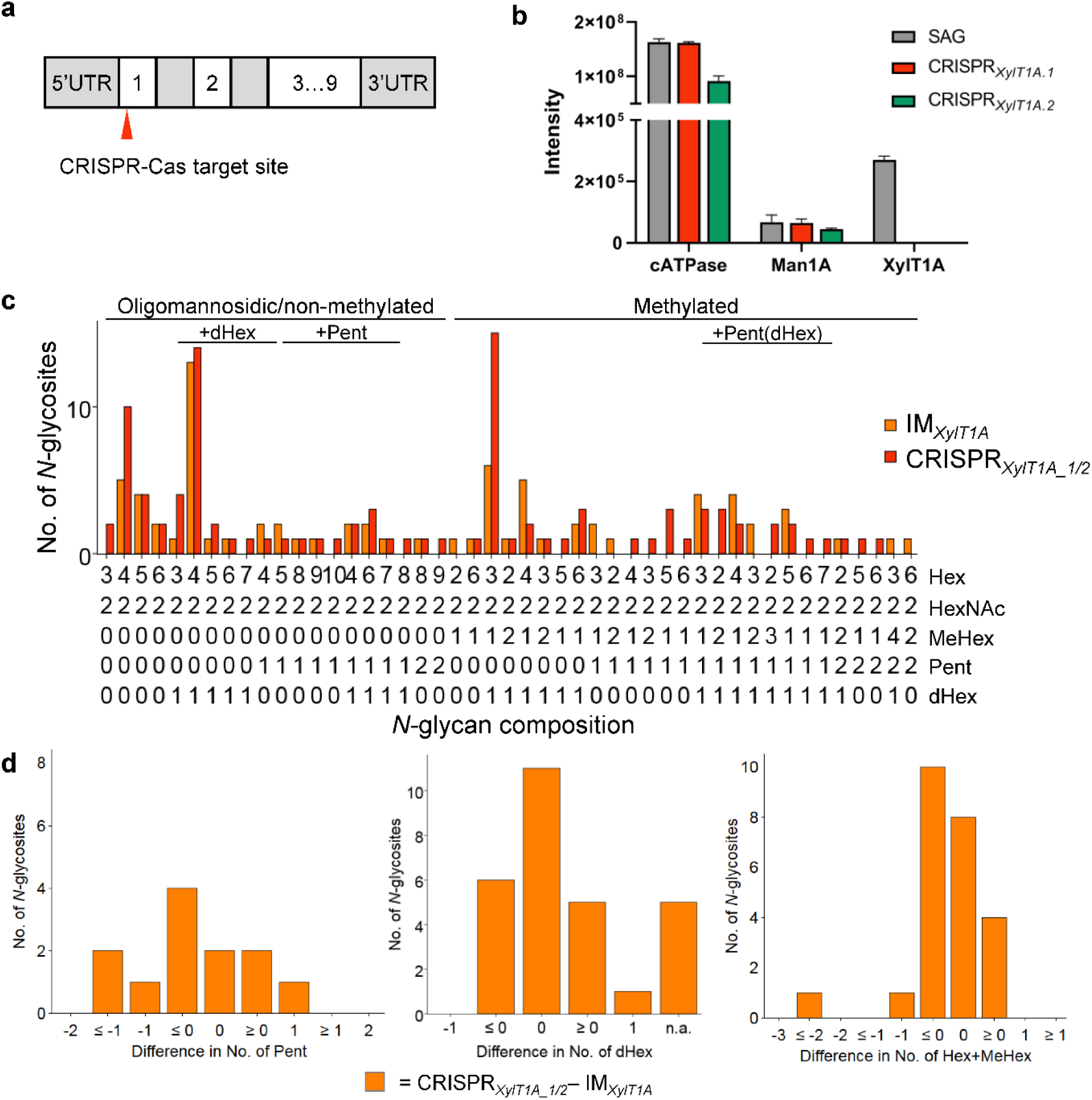
Xylosyltransferase 1A mutant generated via CRISPR/Cas9 supports findings in IM_*XylT1A*_. a, Schematic representation of the XylT1A gene including the site targeted by CIRSPR/Cas9. b, Parallel reaction monitoring was employed to prove the knock out of XylT1A on proteomic level in two mutants generated by CRISPR/Cas9. c, Comparison of *N*-glycan patterns between the XylT1A mutant strains in different genetic backgrounds (IM strain and CRISPR-Cas generated mutants) considering comparable *N*-glycosites. d, Detailed comparison of the *XylT1A* strains in different genetic background looking specifically at Pent (left), dHex (middle) and *N*-glycan length (number of Hex+MeHex, right).

**Figure 4 – Supplementary Figure 2.**
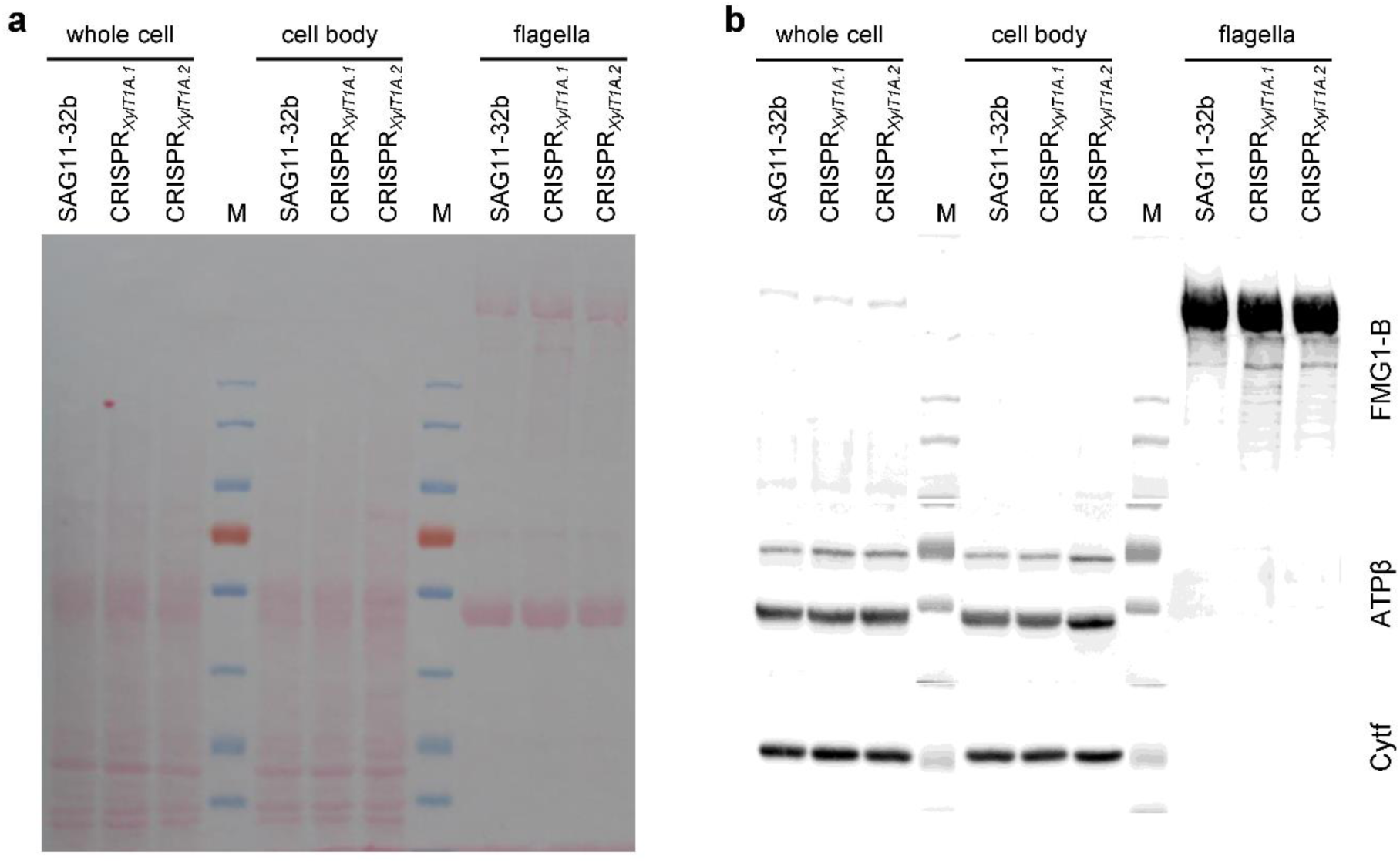
Immunoblot proving the presence of FMG-1B in flagella of CRISPR_*XylT1A.1*_ and CRISPR_*XylT1A.2*_. 30 µg of protein per sample were separated by SDS-PAGE and transferred to a nitrocellulose membrane in biological quadruplicates. a, Ponceau staining of the membrane reveals equal loading between different strains and differential composition of different sample types. b, On one hand, the FMG-1B protein backbone specific antibody proves a correct targeting of FMG-1B to flagella in the mutants despite altered N-glycosylation. On the other, the application of antibodies specifically binding to a chloroplast (Cytf, cytochrome f) marker protein as well as to the ATPase beta subunit (mitochondrial and chloroplast) proves the purity of flagellar samples analyzed.

